# Characterization of bovine vaginal microbiota and its relationship with host fertility, health, and production

**DOI:** 10.1101/2023.12.13.571514

**Authors:** L Brulin, S Ducrocq, G Even, MP Sanchez, S Martel, S Merlin, C Audebert, P Croiseau, J Estellé

## Abstract

**Background:** Because of its potential influence on the host’s phenotype, increasing attention is paid to organ-specific microbiota in several animal species, including cattle. However, ecosystems other than those related to the digestive tract remain largely understudied. In particular, little is known about the vaginal microbiota of ruminants despite the importance of the reproductive functions of cows in a livestock context, where fertility disorders represent one of the primary reasons for culling.

**Results:** In the present study, we aimed at better characterizing the vaginal microbiota of dairy cows through 16S rRNA sequencing, using a large cohort of Holstein cows from Northern France. Our results allowed to define a core microbiota of the dairy cows’ vagina, and highlighted that 90% of the sequences belonged to the Firmicutes, the Proteobacteria, and the Bacteroidetes phyla. The core microbiota was composed of four phyla, 16 families, 14 genera and only one amplicon sequence variant (ASV), supporting the idea of the high diversity of vaginal microbiota within the studied population. This variability was partly explained by various environmental factors such as the herd, the sampling season, the lactation rank and the lactation stage. In addition, we investigated potential associations between the diversity and the composition of the vaginal microbiota and several health-, performance-, and fertility-related phenotypes. Our analyses highlighted significant associations between the α and β- diversities and several traits including the first insemination outcome, the productive longevity, and the culling. Besides, relevant phenotypes were correlated with the abundance of several genera, some of which, such as *Leptotrichia*, *Streptobacillus*, *Methylobacterium-Methylorubrum*, or *Negativibacillus*, were linked to multiple traits.

**Conclusion:** Considering the large number of samples, which were collected in commercial farms, and the diversity of the phenotypes considered, this study represents a first step towards a better understanding of the close relationship between the vaginal and the dairy cow’s phenotypes.

## Background

Microbiotas are complex communities of microorganisms present in a specific habitat. They are gaining interest because of their tight interactions with the host or the environment they inhabit. In livestock science, studies mainly target the associations between the microbiota of a specific organ and the host phenotypes, permitting a better understanding of some host-microbiota interactions [1] which could lead to performance prediction for breeding [2, 3].

In cattle, the rumen microbiota is of particular interest because of the major role the microorganisms play in the ruminal digestion, impacting production, feed efficiency, or methane emission [4–6]. However, the reproductive tract is also of great importance for the livestock sector, reproduction being a cornerstone of dairy farms productivity. Indeed, the focus of selection on milk production has led to a decline of dairy cattle fertility [7, 8], especially because of the negative phenotypic and genetic correlations between reproductive and production traits and the poor heritabilities of fertility phenotypes [9]. These reproductive issues led to increased calving interval and culling rate. Pinedo *et al.* [10] indicated that 17.7% of the animals from 38 states of the USA were culled because of reproduction issues. In Canada, a similar proportion of 15.5% has been reported [11].

The vaginal microbiota of humans [12] and diverse livestock species such as mares [13], sows [14, 15] or ewes [16] has been characterized with the objective to decipher its relationship with fertility impairs [17] or vaginal infections [14]. Despite being underexplored, cattle were found to present a vaginal microbiota with a larger diversity than other species [18]. In addition, there have been studies aimed at uncovering correlations with fertility traits [19, 20]. However, the α and β-diversities in the vagina of the cows which became pregnant after the artificial inseminations (AI) were not significantly different from the ones which failed, and differences were only observed with differentially abundant operational taxonomic units (OTU) [19, 20]. Similar observations were made within the ovine vagina [16].

The bacterial community of the reproductive tract has also been explored because of the negative impacts of some taxa on the animal’s welfare, production performances and fertility through uterine diseases, such as endometritis, metritis or pyometras [21]. *Trueperella pyogenes, Escherichia coli, Prevotella melaninogenicus* and *Fusobacterium necrofurom* have already been identified as pathogens which caused the inflammation or increased the severity of the infection with culture method [22–24]. *Fusobacterium, Bacteroides, Helcococcus* and *Porphyromonas* were also linked with the infectious status according to differential abundance analyses with 16S rRNA sequencing [24, 25]. In addition, a lower α-diversity was generally observed in the infected reproductive tract [24, 25].

However, the aforementioned studies have been performed on a relatively small number of animals observed in experimental conditions, calling for more extensive work in commercial production settings. To this aim, in the present study, we investigated the vaginal microbiota of Holstein cows and heifers and its relationships with health, fertility, and production performance of the host animal. Using a large number of samples collected in French commercial herds, with the 16S rRNA gene amplicon sequencing, we characterized the vaginal microbiota composition of Holstein cows. We also analyzed its variations in terms of α and β- diversities between different physiological and environmental conditions (herd, parity, season) and the traits of interest. Finally, we looked for genera with varying abundance patterns according to the host’s phenotypes.

## Methods

### Animal sampling and phenotyping

A total of 1 171 samples were collected throughout 19 commercial farms located in Northern France, with an average of 51 sampled cows per herd (min = 23, max = 108). Among these samples, 281 were samples from heifers and 890 were collected on cows (parity 1 to 5). Sample collection was conducted between September 2017 and December 2018 by animal reproduction technicians from the Gènes Diffusion company by performing vaginal swabs on non-gestating Holstein cows. The samples were then stored in sterile tubes at -20°C in the Gènes Diffusion research laboratory (Institut Pasteur de Lille, France) until the DNA extraction.

Phenotypic information related to health, production, and reproduction was extracted from routinely collected data at the farms. Features of each trait are presented in Table 1. The reproductive performance of the animals was assessed through two quantitative traits, namely calving interval (CI), corresponding to the number of days between two consecutive calvings, and the calving to fertilizing AI interval (C-AI_f_), and a binary trait indicating success (1) or failure (0) at the first insemination (FIS). For all fertility traits, only the animals sampled before the first insemination were evaluated. Production-related traits were milk, fat and protein yields, as well as fat and protein contents measured during the first 305 days of lactation (MY, FY, PY, FC, and PC, respectively). Only animals with a complete lactation of at least 300 days and no longer than 600 days were considered. The vaginal health status was also recorded during sampling and phenotyped as the presence or absence of infections (metritis, pyometras and any other infection declared by a veterinarian). Finally, two longevity phenotypes were defined as culling or not at the end of the observed lactation (referred as Culling, 0/1) and length of the productive life (referred as Longevity in days).

**Table 1:** Descriptive statistics of the different phenotypes of interest.

| Phenotypes | Units | Number samples | Number herds | Min. | Mean | Max. | S.D. |
| --- | --- | --- | --- | --- | --- | --- | --- |
| **Calving interval (CI)** | days | 422<br>(338) | 17<br>(17) | 311<br>(311) | 419.6<br>(422.3) | 685<br>(685) | 73.51<br>(75.70) |
| **Calving to fertilizing AI interval (C-AI<sub>f</sub>)** | days | 430<br>(339) | 17<br>(17) | 43<br>(43) | 142.2<br>(141.1) | 466<br>(396) | 77.38<br>(72.98) |
| **Milk yield (MY)** | kg/305d | 545<br>(433) | 16<br>(16) | 3 530<br>(3 530) | 9 072<br>(9 026) | 14 275<br>(14 275) | 1 792.37<br>(1 793.60) |
| **Protein content (PC)** | g/kg/305d | 541<br>(430) | 16<br>(16) | 27.2<br>(27.2) | 31.5<br>(31.4) | 37.2<br>(37.0) | 2.00<br>(2.06) |
| **Protein yield (PY)** | kg/305d | 543<br>(432) | 16<br>(16) | 1 428<br>(1 428) | 2 847<br>(2 831) | 4 262<br>(4 262) | 555.68<br>(554.57) |
| **Fat content (FC)** | g/kg/305d | 529<br>(428) | 16<br>(16) | 29.2<br>(29.6) | 38.9<br>(39.2) | 49.9<br>(52.6) | 4.07<br>(4.29) |
| **Fat yield (FY)** | kg/305d | 543<br>(431) | 16<br>(16) | 1 751<br>(1 751) | 3 531<br>(3 520) | 5 896<br>(5 889) | 755.12<br>(749.83) |
| **Longevity** | days | 402<br>(336) | 16<br>(16) | 157<br>(157) | 1 258<br>(1 269) | 2 562<br>(2 562) | 543.35<br>(535.06) |
| **Culling** | Qualitative<br>(0/1) | 522<br>(410) | 16<br>(14) | - | - | - | - |
| **First insemination success (FIS)** | Qualitative<br>(0/1) | 386<br>(302) | 13<br>(12) | - | - | - | - |
| **Infection** | Qualitative<br>(0/1) | 249<br>(167) | 6<br>(5) | - | - | - | - |

The numbers in brackets refer to the rarefied dataset. Longevity = length of dairy career from first calving to the end of the last lactation, Culling = animal not cull/cull at the end of the lactation (0/1), Infection = absence/presence of a reproductive tract infection at the sampling date (0/1).

### Microbiota DNA extraction

The DNA extraction was performed using the Nucleospin® 96 Soil kit (Macherey Nagel) under aseptic conditions at room temperature and following manufacturer’s recommendations. In brief, the cotton swabs used for sampling were first cut and transferred to 2 mL tubes where supplied ceramic beads were then transferred. Lysis buffer and Enhancer buffer were added to tubes and they were agitated at 30 Hz for 2 min with homogenizer (Tissue Lyzer) to break and lyse the samples. The protocol followed the supplier’s recommendations until the elution phase of the samples with 50 µL of TE 1X pH 8.0 preheated at 70 °C followed by a 1-hour incubation at room temperature. A centrifugation at 6000 x g for 2 min was performed and the DNA was stored at -20°C until use.

### Library preparation and 16S sequencing

The sequencing library is based on a dual-indexed paired-end sequencing strategy targeting the V3-V4 variable regions of the 16S rRNA gene, as previously described [26]. In brief, to achieve this, two PCRs were successively applied: from 2 µL of the extracted DNA diluted to 1/25, a first PCR was realized in a final volume of 50 µL, using 2.5 U of Precision Taq Polymerase (Applied Biological Materials Inc, Richmond), each primer had a final concentration of 500 nM. For this first PCR, forward and reverse primers had been designed with the 5’-Tag sequences 5’-TCGTCGGCAGCGTCAGATGTGTATAAGAGACAG-3’ and 5’- GTCTCGTGGGCTCGGAGATGTGTATAAGAGACAG-3’ for the forward and reverse primers, respectively, and 16S rRNA gene specific sequences 5’-CCTACGGGNGGCWGCAG-3’ and 5’- GACTACHVGGGTATCTAATCC-3’ for forward and reverse primers, respectively. According to *Escherichia coli* 16S rRNA sequence gene specific primers target a locus between position 341 and 785, resulting in the amplification of a locus of 445 bp. The amplification conditions were 3 min at 94 °C, 25 cycles of 15 s at 94 °C for denaturation, 15 s at 51 °C for primers annealing and 45 s at 68 °C for extension, followed by an incubation at 68 °C for 1 min. At the end of this first PCR, amplification products had been purified with NucleoFast® 96 PCR (Macherey Nagel) according to supplier recommendations except for the last step for which 30 µL of TE 1X pH 8.0 preheated at 70 °C had been used for elution. From 5 µL of the previously purified PCR products, a second PCR was performed in a final volume of 50 µL, 2.5 U of Precision Taq Polymerase (Applied Biological Materials Inc, Richmond). Each primer had a final concentration of 500 nM. The amplification conditions were the same as those of the previous one except the number of cycles reduced to 8. In addition to the Tag sequences, these PCR2 primers contain a locus to index the samples (barcode sequence) and a locus sequence adapter suitable for the Illumina sequencing technology. A NucleoFast® purification step identical to the one presented above was performed followed by a Quant-iT PicoGreen ds DNA quantification (Life Technologies). An equimolar pool of the library was produced, 200 µL of this mixture was purified by NucleomagNGS® (Macherey Nagel). The purification was performed twice to 1.2X with 240 µL of beads suspension at each purification to conclude with a final elution in 50 µL of TE 1X pH 8.0. This purified pool library was then monitored by Bioanalyzer with the High sensitivity DNA Chips kit (Agilent) as a quality control and to estimate the average size of the pool. Then, a quantification of DNA was realized by Qubit assay (Invitrogen). The concentration of DNA and the average size were used to assess molarity of the purified pool.

Sequencing library has been paired-end sequenced on Miseq platform (Illumina) with MiSeq Reagent Kit v3 allowing 600 sequencing cycles to be performed. At the end of the sequencing a quality control by FastQC v.0.11.9 [27] was carried out.

### Bioinformatic processing of 16S data and taxonomic assignment

The sequence analyses were conducted using the R software (v.4.2.1) [28] and the dada2 v.1.24-0 package [29] following the author’s recommendations. Each sequencing run was analyzed separately for the quality filtering, denoising pair-end merging, and amplicon variant calling steps. Primers and indexes were trimmed and the forward and reverse reads were truncated using the Phred score Q20 as quality threshold [30]. Thus, forwards reads were trimmed at positions between 280 bp and 290 bp and reverse reads were trimmed at positions between 220 bp and 230 bp. The DADA2 method [29] was chosen to cluster the sequences with a pairwise identity threshold of 100% (Amplicon Sequence Variants - ASV) and to obtain a count matrix of samples by ASV. All lab batch-specific tables were merged into a unique count table with chimeras removed.

The SILVA v.138 [31] was used for taxonomic assignment of the ASVs at all taxonomic ranks, from reign to the species level. To avoid sequence depth bias in diversity analyses, each sample was rarefied to a common depth of 7 000 sequences by using the phyloseq (v.1.40.0) R package [32].

### Statistical analysis

α-diversity was analyzed on the rarefied dataset as the observed richness and the Shannon diversity index using the phyloseq and vegan (v.2.6-2) [33] R packages, respectively. Factors significantly associated with α-diversity were identified with analyses of variance (ANOVA) of the car (v.3.1-0) [34] package. This analysis was performed to identify the co-factors significantly associated with α-diversity, the latter being set as the response variable. Then, associations between richness or Shannon index and the quantitative traits in productive cows (parity ≥ 1) were explored with linear models. First, linear models were performed using the α-diversity metrics and the phenotypes as the dependent variables and all factors, previously found with significant effects, as independent variables. Finally, Pearson’s correlation was calculated using the residuals of α-diversity and of the phenotypes to test for association.

β-diversity was also computed using the Bray-Curtis dissimilarity matrix, computed on the rarefied dataset with the vegan R package. Bray-Curtis values were used in a permutational multivariate analysis of variance (PERMANOVA) to evaluate the potential associations between the β-diversity and the phenotypes of interest in productive cows (parity ≥ 1). The herd, the lab batch, the season of sampling, the days in milk (DIM), the parity and the phenotype were considered as cofactors. The marginal effects of each factor were assessed using the by = ”margin” option.

Finally, differential abundance analyses at the genus taxonomic level on the unrarefied table were performed using the analysis of compositions of microbiomes with bias correction (ANCOM-BC) method using the ANCOMBC R package (v.1.4.0) [35]. Taxa observed in less than 90% of samples were removed as well as samples with less than 1,000 sequences. The p-values were adjusted using the Benjamini-Hochberg procedure. The level of significance was set at p ≤ 0.05 for all statistical analyses and tendencies were defined as being 0.05 < p ≤ 0.1.

## Results

### Taxonomic composition of the bovine vaginal microbiota

Sequence analyses revealed 37 840 ASVs within the vaginal microbiota of the entire population of cows and heifers, representing an average of 439 ASVs per sample. We observed 30 153 ASVs considering the cows only, with an average of 473 ASVs per sample. This figure was of 29 631 ASVs when limiting the analysis to the healthy cows, with an average of 478 ASVs per sample. In heifers, 18 203 ASVs had been detected, with an average of 341 ASVs per sample.

A total of 33 phyla were detected among all the healthy animals (Figure 1-A). Overall, Firmicutes, Proteobacteria and Bacteroidetes were the most prevalent phyla, representing 90% of the reads, with a relative abundance of 42%, 36% and 12%, respectively. At the genus level, 17 genera had a relative abundance superior to 1%: *Escherichia-Shigella* (16.9%), *Photobacterium* (7.5%), *UCG-005* (6.9%), unknown genus from UCG-010 family (6.9%), *Histophilus* (4.3%), *Ureaplasma* (3.8%), *Rikenellaceae RC9 gut group* (3.5%), an unknown genus from Oscillospirales family (3.1%), *Bacteroides* (2.8%), an unknown genus from Pasteurellaceae family (2.1%), *Alistipes* (1.8%), unknown genus from Lachnospiraceae family (1.7%), unknown genus from Bacteroidales RF16 group family (1.5%), *Prevotellaceae UCG-003* (1.3%), *Phyllobacterium* (1.2%), *Romboutsia* (1.2%) and *Monoglobus* (1.2%).

**Figure 1:**
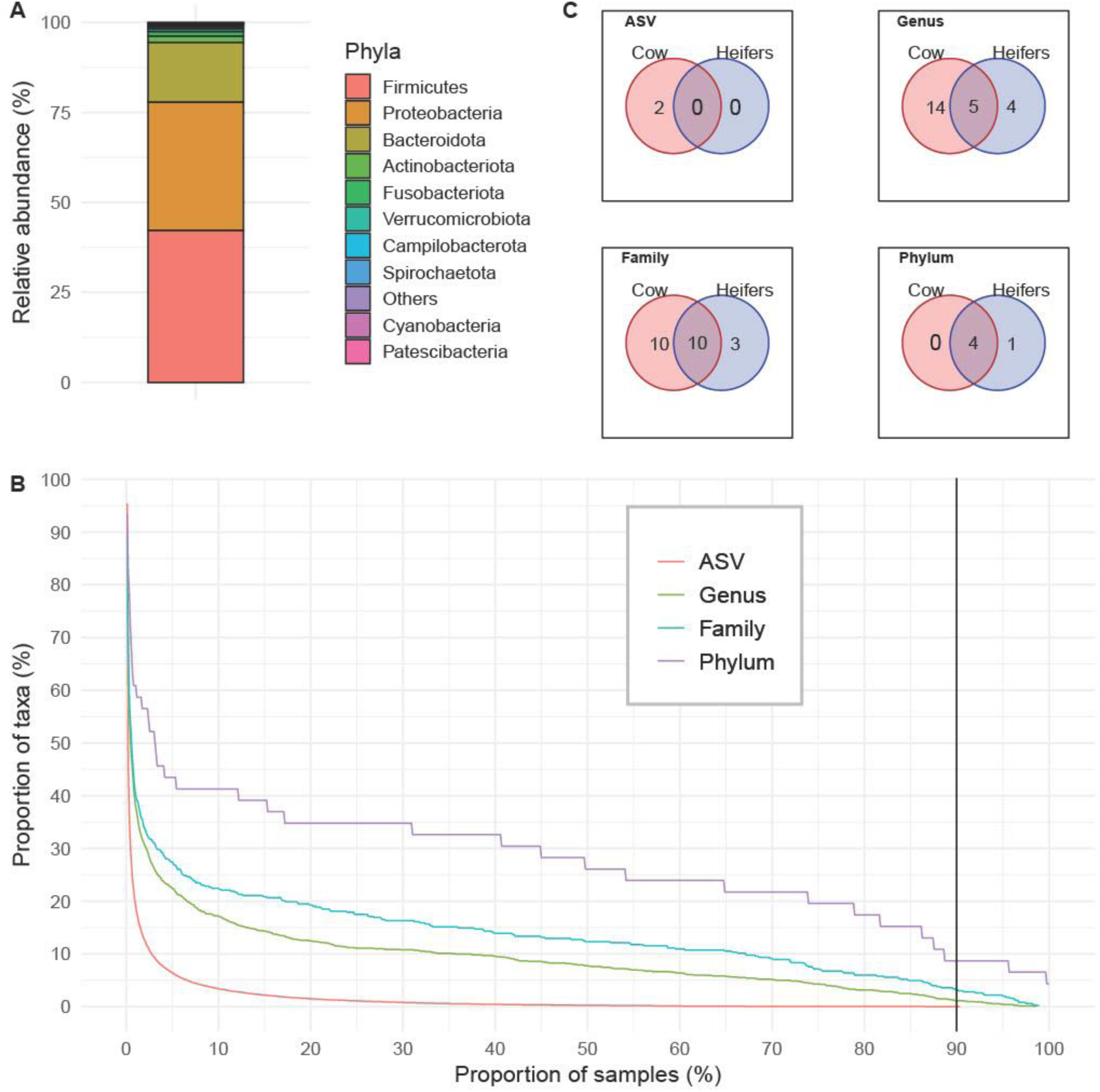
Characterization of the bovine vaginal microbiota. A. Relative abundance of the 10 most abundant phyla in the vagina. The other observed phyla are included in the “Others” category; B. Proportions of taxa shared by the samples for each taxonomic rank. The vertical line represents the minimum threshold (90%) for the taxa to belong to the microbiota core; C. Number of taxa shared by the core microbiotas of the cows and heifers at different taxonomic ranks.

We investigated the composition of the vaginal core microbiota using ASVs, genera, families and phyla that were present in at least 90% of the healthy animals. This relaxed threshold was chosen because of the large number of samples and the various environmental conditions in our study. Only one ASV, 14 genera, 16 families and 4 phyla were commonly found in vaginal microbiota, which represented a proportion of <0.0001%, 1.5%, 5.2% and 12.1%, respectively. Conversely, as highlighted with the sharp decrease in Figure 1-B, half of the ASVs, genera, and families were shared by less than 3% of the vaginal samples.

The core vaginal microbiota of cows and heifers exhibited some differences (Figure 1 - C) with five genera, 10 families and four phyla shared between the two groups and 18 genera, 13 families, and one phylum specifically observed in either cows or heifers. Interestingly, only two core ASVs, related to *UCG-005* (Oscillospiraceae family) and *Romboutsia* (Peptostreptococcaceae family), were found in the cows’ samples.

### Links between vaginal microbiota diversity and possible cofactors

The α-diversity of the whole dataset, measured as the Shannon index, was 4.404 and 2.919 for the ASV and the genus taxonomic ranks, respectively. The samples presented an average Bray-Curtis distance value of 0.801 at the ASV level and of 0.621 at the genus level. More specifically, lactating cows had an average Shannon index of 4.545 and 2.980 at the ASV and genus ranks, respectively. Besides, they presented an average Bray-Curtis distance of 0.793 for the ASV and 0.599 for the genus.

All the tested physiological or environmental factors presented a significant association with the α and β-diversities of the bovine vaginal microbiota (Table 2), both considering ASVs and genera. Interestingly, the α-diversity was found to increase with DIM and parity meaning that older animals and animals which had longer intervals between calving and sampling had a higher α-diversity.

**Table 2:** Associations between cofactors, phenotypes and the α and β-diversities for ASV and genus taxonomic ranks. Bold p-values indicate significant p-values (p ≤ 0.05).

|  |  | Observed richness |  | Shannon |  | Beta-diversity |  |
| --- | --- | --- | --- | --- | --- | --- | --- |
|  | Samples | ASV | Genus | ASV | Genus | ASV | Genus |
| Factors |  |  |  |  |  |  |  |
| Lab batch | 823 | < 2.2e-16 | 8.042e-09 | 2.258e-13 | 2.708e-13 | 0.0001 | 0.0001 |
| Herd | 823 | 3.204e-12 | 4.003e-12 | 1.654e-11 | 1.404e-10 | 0.0001 | 0.0001 |
| Season | 823 | 0.0001 | 0.009 | 0.0005 | 0.007 | 0.0001 | 0.0001 |
| Lactation rank | 823 | 0.002 | 0.029 | 0.0001 | 0.018 | 0.0001 | 0.0001 |
| Days in milk (DIM) | 823 | 0.004 | 0.005 | 1.565e-05 | 0.0001 | 0.0002 | 0.0001 |
| Phenotypes |  |  |  |  |  |  |  |
| Culling | 410 | 0.0006 | 0.002 | 0.010 | 0.010 | 0.014 | 0.019 |
| Longevity | 336 | 0.137 | 0.305 | 0.421 | 0.487 | 0.014 | 0.019 |
| Infection | 167 | 0.356 | 0.280 | 0.249 | 0.164 | 0.191 | 0.191 |
| CI | 338 | 0.357 | 0.093 | 0.167 | 0.147 | 0.178 | 0.205 |
| C-Al<sub>f</sub> | 339 | 0.180 | 0.072 | 0.107 | 0.119 | 0.183 | 0.174 |
| FIS | 302 | 0.047 | 0.035 | 0.018 | 0.034 | 0.040 | 0.014 |
| MY | 433 | 0.984 | 0.728 | 0.560 | 0.822 | 0.225 | 0.228 |
| PY | 432 | 0.607 | 0.742 | 0.215 | 0.269 | 0.167 | 0.204 |
| FY | 431 | 0.644 | 0.716 | 0.253 | 0.178 | 0.223 | 0.365 |
| PC | 430 | 0.239 | 0.141 | 0.200 | 0.094 | 0.461 | 0.586 |
| FC | 428 | 0.588 | 0.333 | 0.355 | 0.108 | 0.186 | 0.264 |

The significantly associated β-diversity, expressed with the Bray-Curtis distance, in PERMANOVA analyses highlighted the variability of the community structure that was associated with these factors representing 38.1% of the total variance. More specifically, herd, parity, season and DIM were associated with explained variances of 6.0%, 1.6%, 0.72% and 0.37%, respectively.

### Associations between the vaginal microbiota and the host’s longevity

Both the α and β-diversities were significantly associated with the decision of ending the production life (that is, the culling) at the end of the lactation for both taxonomic ranks, with the cull animals having higher α-diversity. Conversely, β-diversity was significantly associated only with the length of the productive longevity (Table 2).

The differential abundance analysis pointed out 47 genera which were more frequently observed in cull animals (Figure 2 – A), with the *Pseudomonas* (0.22%) genus being the most abundant in cull animals. In contrast, only one genus, *Negativibacillus* (0.09%), was more abundant in animals that entered a subsequent lactation. *Negativibacillus* was also among the six genera that were commonly observed in long-career animals with *Ruminobacter* (0.17%), *Negativibacillus* (0.09%) (also less abundant in cull animals), *Parasutterella* (0.16%), *Anaerovibrio* (0.03%) and two unknown genera from *Paludibacteraceae* (0.57%) and *Peptococcaceae* (0.18%) families (Figure 2-B). *Methylobacterium-Methylorubrum* (0.06%) was the sole genus negatively associated with longevity (and was also overabundant in cull animals).

**Figure 2:**
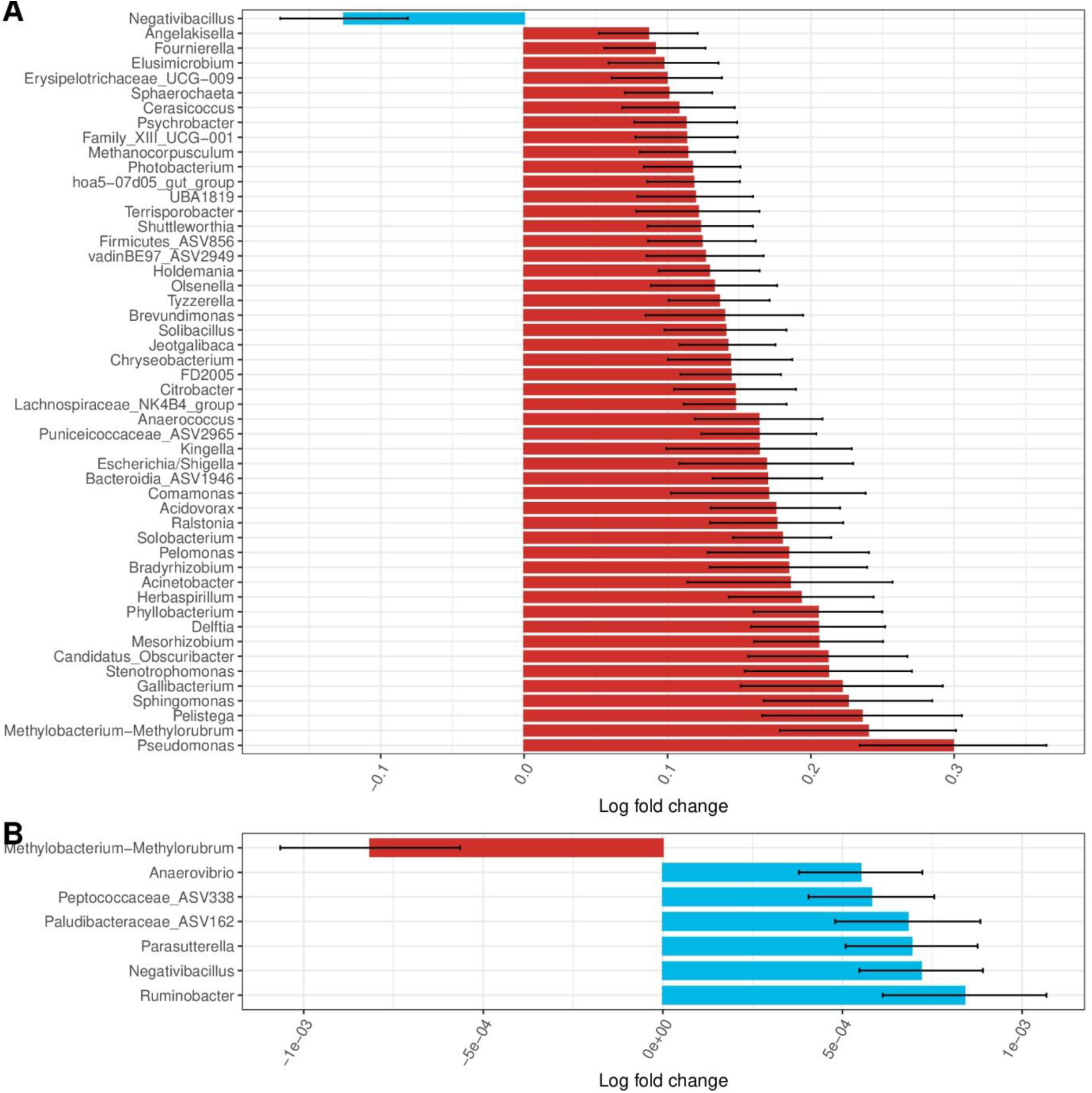
Log-fold change values of the genera associated with the Culling and Longevity. A. Culling = animal cull or not at the end of the lactation (0/1), B. Longevity = length of dairy career from first calving to the end of the last lactation. The red bar genera were associated with the cull or short-career animals whereas the blue bar genera more abundant in long-career animals or in those that were not culled at the end of the lactation.

### Associations between the vaginal microbiota and the host’s vaginal health

We hypothesized that the microbiota of infected vaginas would be significantly different from the healthy vaginal microbiota. Even if the infectious status of the bovine vagina was not correlated with the α and β-diversities index at the ASV level or at the genus taxonomic rank (Table2), the differential abundance analysis pointed out significant associations between the vaginal infection status and 52 genera (Figure 3). Pathogenic genera such as *Peptoniphilus* (0.08%), *Porphyromonas* (0.51%) and *Fusobacterium* (0.28%) were found overabundant in infected vaginas whereas commensal genera, as *Streptobacillus* (0.59%) and *Leptotrichia* (0.16%) were less abundant in healthy bovine vagina. *Campylobacter* (0.51%) genus, represented by a unique ASV assigned to the *C. lanienae* species (0.03%), was also strongly and positively associated with a healthy vagina.

**Figure 3:**
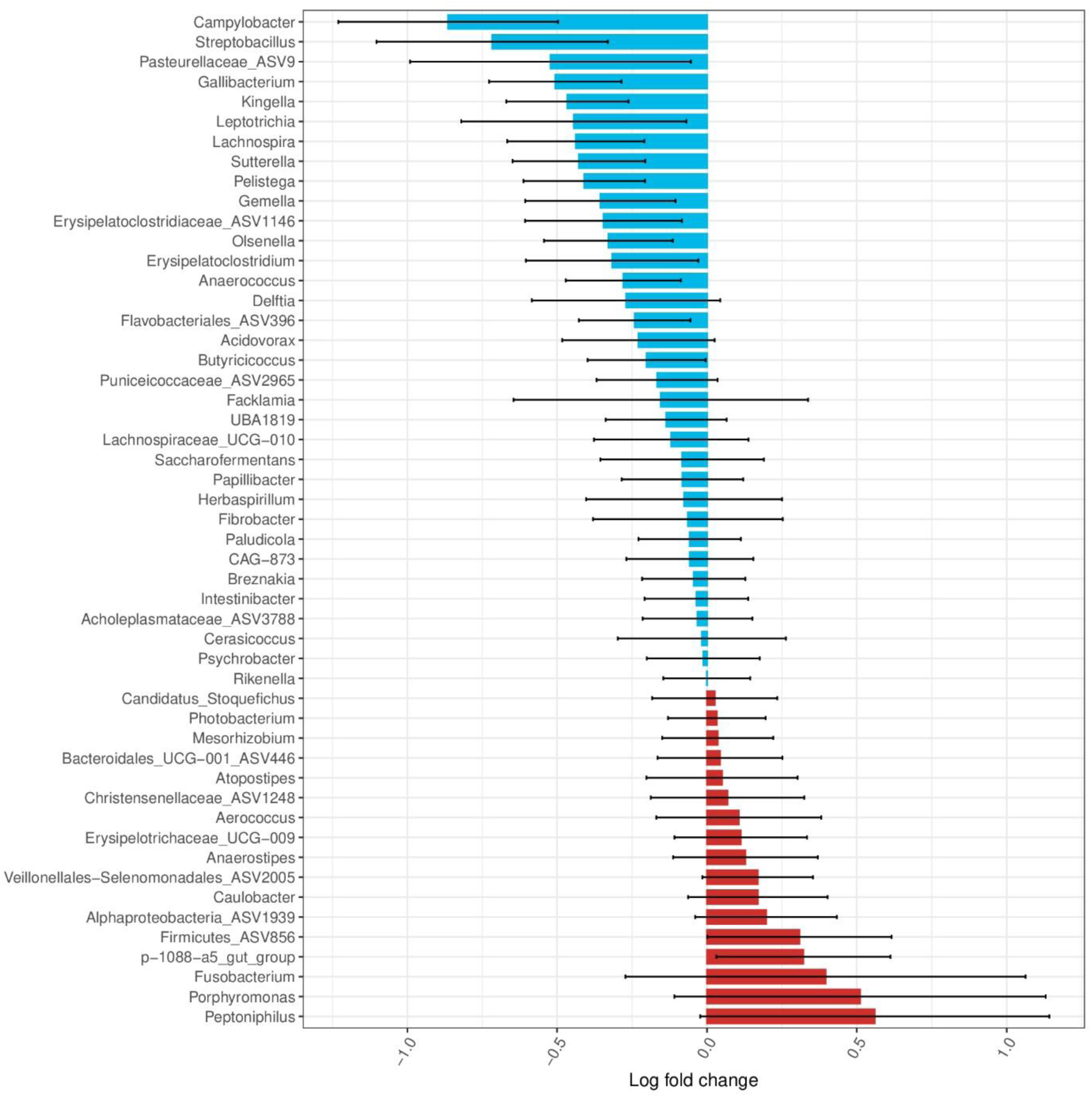
Log-fold change values of the genera associated with the health status of the animals. The red bar genera were associated with infected vaginas whereas the blue bar genera were associated with healthy vaginas.

### Associations between the vaginal microbiota and the host’s fertility traits

The analyses did not show significant associations between the α and β-diversities and the quantitative fertility traits CI and C-AI_f_ (Table 2). However, FIS presented significant associations with both the observed richness and the Shannon index. In this sense, animals that were not pregnant after the first AI, generally had an increased α-diversity score. Furthermore, the β-diversity was also significantly associated with the FIS trait, a result supported by the differential abundance analysis. More specifically, *Streptobacillus* (0.59%) and *Methanosphaera* (0.004%) were both significantly more abundant in cows pregnant after a unique AI (Figure 4 – A). Interestingly, CI and C-AI_f_ interval traits were also negatively associated with a few genera (Figure 4 – B and C). Among these, *Leptotrichia* (0.16%) and *Fournierella* (0.007%) were both more abundant in animals with shorter CI and C-AI_f_ intervals.

**Figure 4:**
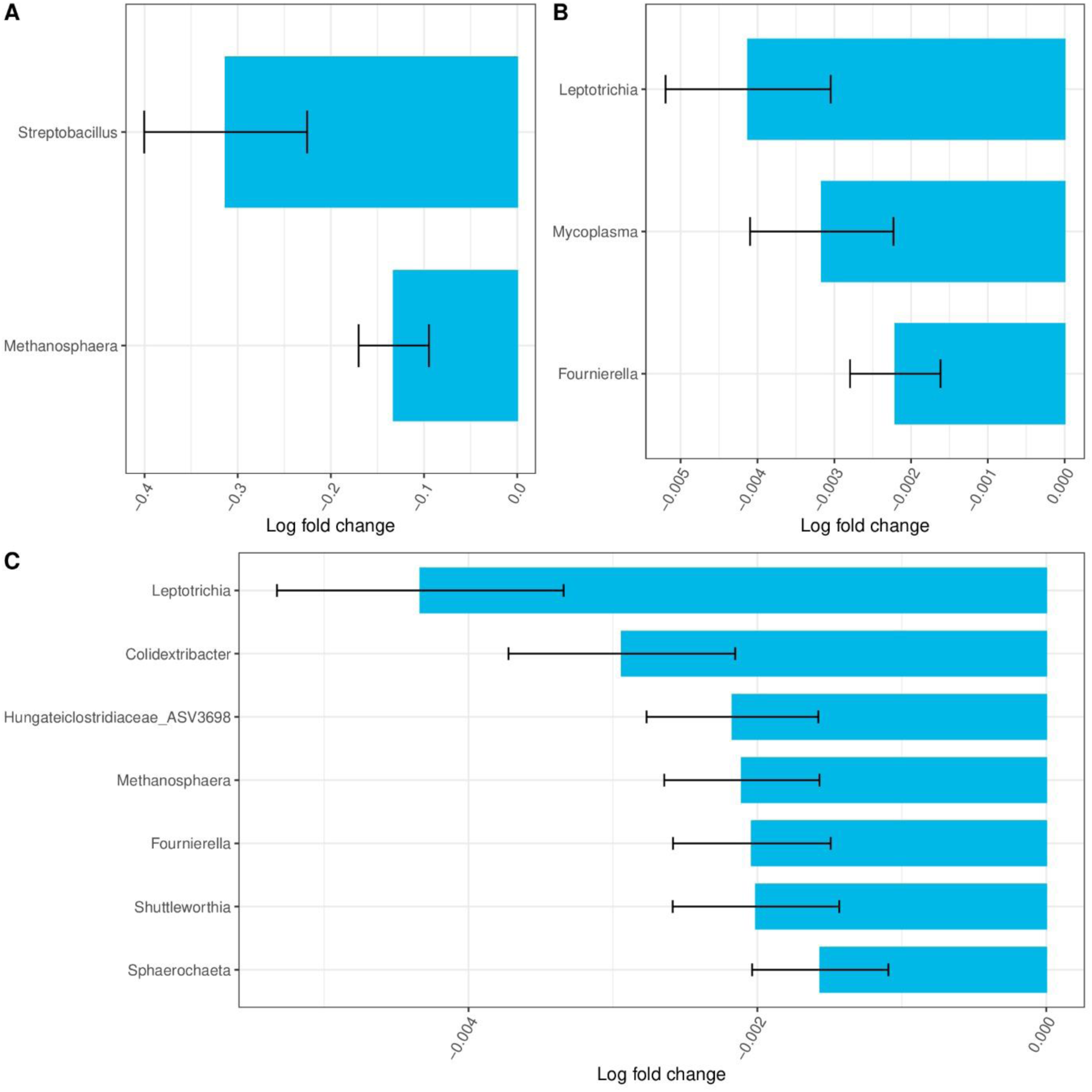
Log-fold change values of the genera associated with fertility traits. A. FIS = first insemination outcome (0/1), B. CI = calving interval, C. C-AI_f_ = calving to fertilizing AI interval. The red bar genera were more abundant in animals with poorer fertility records whereas the blue bar genera were more abundant in animals with interesting fertility results.

### Associations between the vaginal microbiota and the host’s production

None of the production-related traits evaluated in our study was significantly associated with change in α or β-diversities in the vaginal microbiota (Table 2). Thus, the richness of the number of taxa or the overall ecosystem structure of the vaginal microbiota does not seem to change with the animal dairy performances.

Yet, considering the differential abundance analysis, multiple genera were differentially abundant depending on the lactation performance of the animals (Figure 5). In this sense, 21 genera were significantly more abundant in the vagina of animals with higher milk, protein and fat yields (Figure 5 – A). Among these genera, *Lachnospiraceae AC2044 group* (0.09%) was the genus which had the strongest associations with the three phenotypes. Besides, *Ruminobacter* (0.17%) was also among the ten most significant genera for the three traits. In contrast, 20 other genera were significantly more abundant in animals that had reduced performances for the milk, fat, and protein yields (Figure 5 - A). *Streptobacillus* (0.59%), *Histophilus* (4.3%), *Ureaplasma* (3.8%) and *Facklamia* (0.29%) were among the 10 genera which had the strongest associations with reduced performances for the three traits (Figure 5 – B to D). We also looked for significant associations with the fat and protein content in the milk. Only tendencies were highlighted among which *Bifidobacterium* (p = 0.096) (0.35%), *Atopostipes* (p = 0.096) (0.044%), and *Clostridium sensu stricto 1* (p = 0.096) (0.58%) tended to be overabundant in animal with higher fat content.

**Figure 5:**
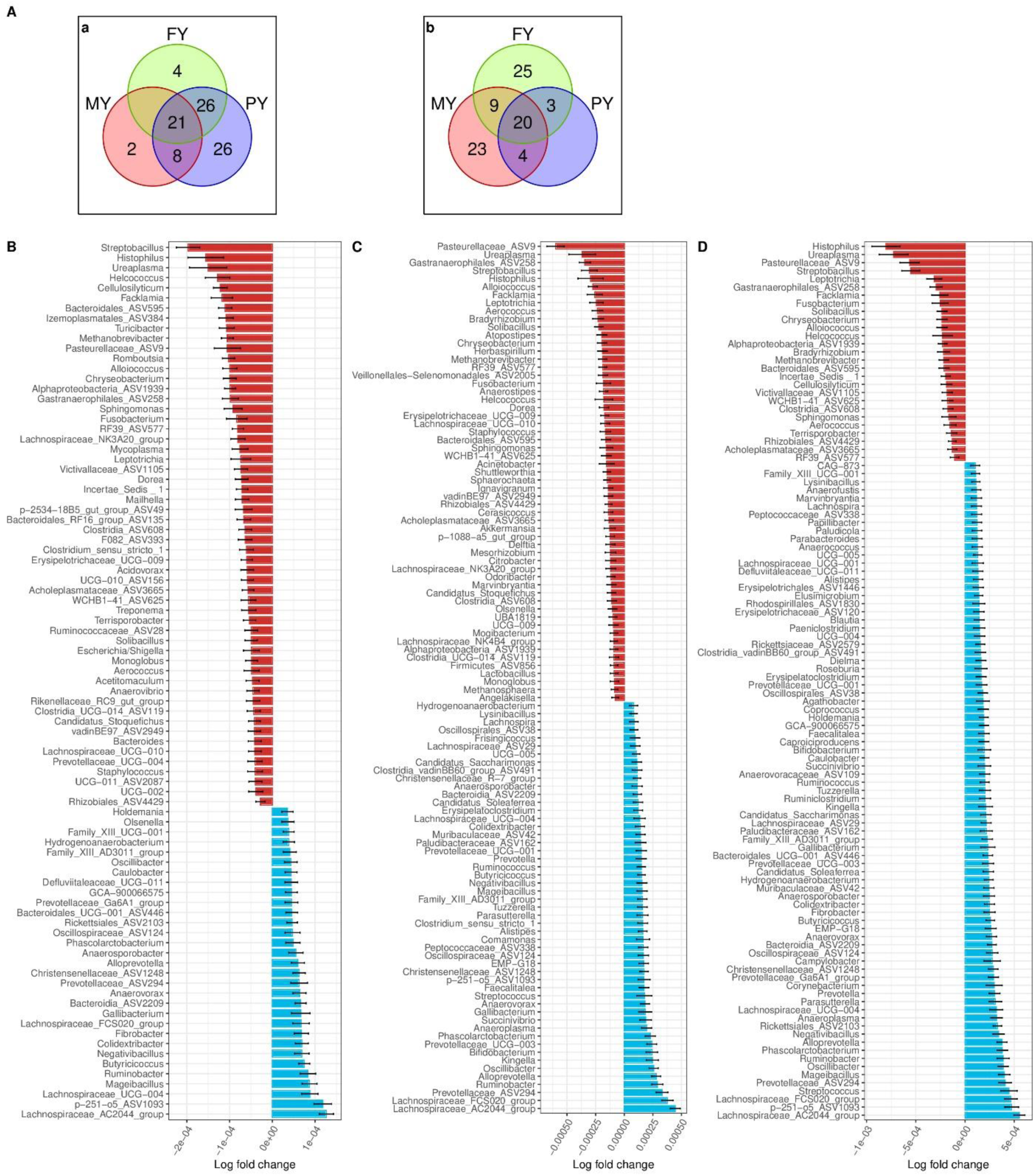
Results of the differential abundance analyses on production traits. A. Number of shared genera between Milk Yield (MY), Protein Yield (PY) and Fat Yield (FY) which were associated with good performances (a) or bad performances (b); B. Log-fold change results of the genera associated with the MY; C. Log-fold change results of the genera associated with the FY; D. Log-fold change results of the genera associated with the PY. For histograms, red bar genera were more abundant in animals with the poorer records whereas the blue bar genera were more abundant in animals with the highest production records.

### Comparative analysis of the genera significantly associated with host traits

The host phenotypes analyzed here were significantly associated with a total of 186 genera composing the vaginal microbiota and 69% were linked to more than one phenotype (Figure 6) with a beneficial and/or a detrimental association. Interestingly, MY, FY and PY were the phenotypes that shared the highest number of genera with same direction correlations (*i.e*. beneficial or detrimental), with a total of 30 genera. The *Negativibacillus* genus was associated in a similar way with the highest number of phenotypes, its abundance being positively associated with the three production traits, the culling, and the longevity.

**Figure 6:**
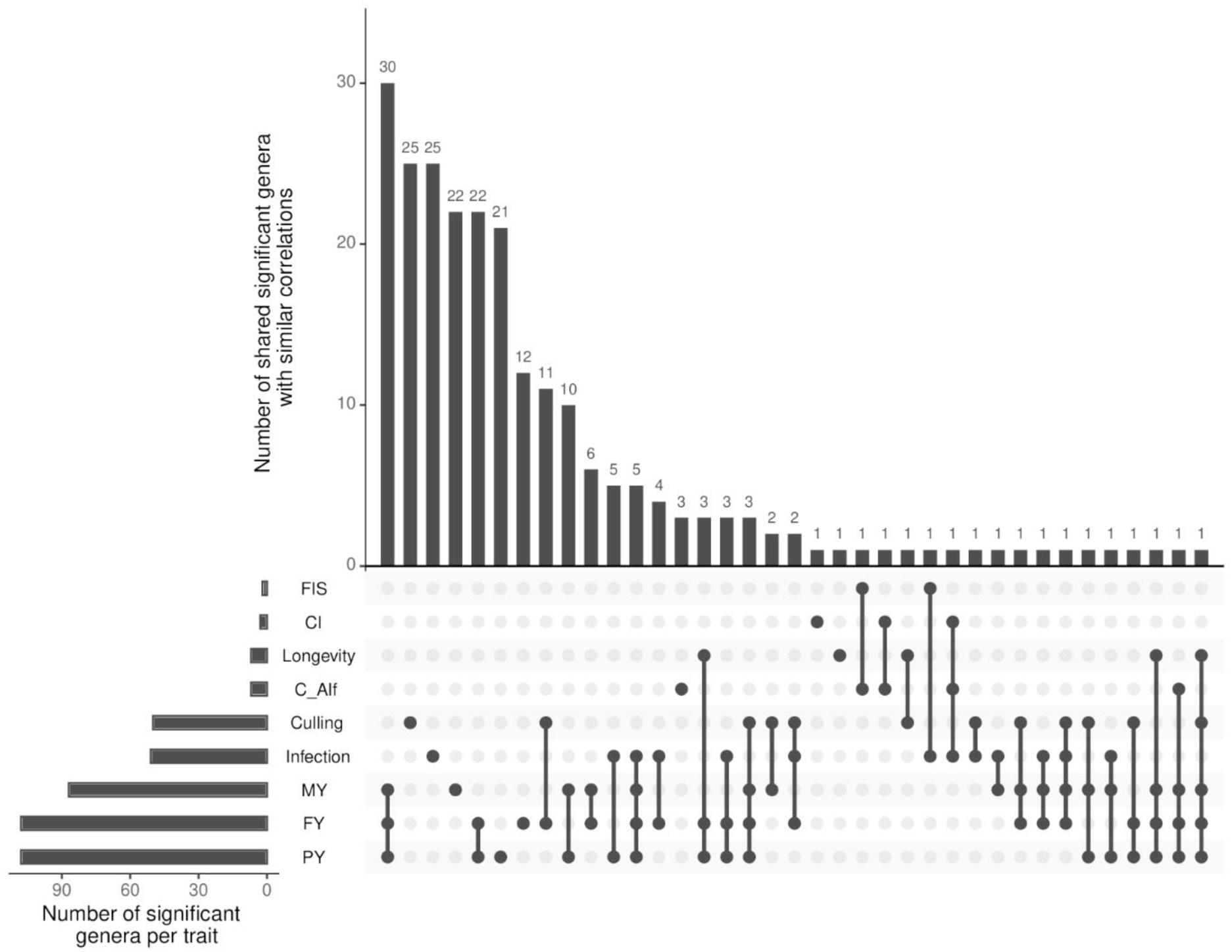
Number of significant genera by trait and shared among traits with same direction relationships. CI = calving interval, C-AI_f_ = calving to fertilizing AI interval, MY = milk yield, PC = protein content, PY = protein yield, FC = fat content, FY = fat yield, Longevity = length of dairy career from first calving to the end of the last lactation, Culling = animal not cull or not at the end of the lactation (0/1), FIS = first insemination outcome (0/1), Infection = absence/presence of a reproductive tract infection at the sampling date (0/1).

Conversely, 32 genera were significantly linked with various phenotypes in a discrepant way. For instance, *Leptotrichia* was more abundant in animals that had shorter CI and C-AI_f_, but was also more abundant in animals with reduced performances in milk, protein and fat yields. Similarly, *Streptobacillus* appeared favorable for FIS and the health status but detrimental regarding production performances.

## Discussion

We present here an unprecedented view of the vaginal microbial communities in dairy cows, which uses, to our knowledge, the largest cohort of animals to date. Our results highlight the complexity of the reproductive tract microbiota, and unveil its links with several traits of major breeding interest. Furthermore, since it is performed on commercial herds, our study is directly representative of current production systems.

### Characterization of the vaginal microbiota of Holstein cows

We showed that the vaginal microbiota is mainly structured into three phyla, namely, *Firmicutes*, *Proteobacteria*, and *Bacteroidetes*. This result is consistent with prior knowledge generated in smaller cohorts in dairy cattle [19, 36], beef [37], and Nellore cattle [38]. In these analyses, *Firmicutes* appeared as a major phylum of the vaginal microbiota with a relative abundance ranging from 32.4% to 65.9%, which is consistent with the relative abundance of 42% found in our study. Interestingly, these phyla are also predominant in other species such as in sows [14, 15] or mares [13]. Our results were more contrasted at the genus level, with 17 genera representing 75% of the taxa, with a maximal relative abundance of 17% for *Escherichia-Shigella* genus. A similar trend was observed by Quereda *et al.* [30] and Clemmons *et al.* [37] who identified 17 and 10 genera with relative abundance ≥ 1% in dairy heifers and cows, respectively. However, only some of the most prevalent genera we identified were also observed by the other authors: only *UCG-005*, *Histophilus*, *Ureaplasma*, *Rikenellaceae R9 group*, *Bacteroides* and *Alistipes* were also predominant in the heifers’ vaginas of Quereda *et al.* [30] and an undetermined genus from *Bacteroidales* order and *Ureaplasma* were also among the most abundant in the heifers of Clemmons *et al.* [37]. These differences could be technical, caused by the sample size, the rearing conditions, or the method used for the analyses (ASV vs OTU).

The high abundance we found for *Escherichia-Shigella* (17%) was unexpected considering its potential implication with metritis occurrence [14, 39], however, this genus was also observed in the vaginal microbiota of beef heifers [20] and other works point *Ureaplasma* [19, 40] as a dominant taxa despite its well-known pathogenicity. These findings illustrate that dairy cows could naturally carry potential pathogens in their reproductive tract. Also, despite its overall abundance, *Escherichia-Shigella* was not considered as part of the core microbiota.

The core microbiota of our vaginal samples was composed of one ASV, 14 genera, 16 families, and four phyla. The definition of the core microbiota could differ across studies, and we choose an approach that aims at providing a balanced estimate in our conditions of sample size and diversity of farms of origin. Quereda *et al.* [30], who define the core microbiota as being present in all samples, have also observed the *UCG-005*, the *Bacteroides* and an unknown genus from the *Ruminococcaeae* in the vaginal core microbiota of dairy heifers. Thus, only a small fraction of the ASVs, genera, families and phyla were shared between animals. More than half of the ASVs, genera and families were present in less than 3% of the samples, indicating that a major part of the taxa could be considered as rare. This finding highlights the diversity of vaginal microbiotas that we could observe in the dairy cows, in agreement with a previous observation made by Miranda-CasoLuengo *et al.* [25].

### The vaginal microbiota is highly environment-dependent

It is generally accepted that microbiota composition and functionalities are highly environment-dependent [41]. Thus, considering the high diversity of taxa we observed, we expected that the vaginal microbiota diversity could be linked to the multiplicity of conditions the cows faced. We were able to evidence associations between both the α and β-diversities and various environmental and physiological factors, including farm management, lab management, season, parity, and lactation stage.

Both the vaginal microbiota diversity and composition were closely related to the herd in the present study. Indeed, the rearing farm integrates numerous variables such as housing conditions, access to grazing land and diet, and has already been found to influence the microbiota of cattle [42, 43]. In addition, within the same herd, other impacting factors were found.

At the animal level, multiparous cows had a higher α-diversity than primiparous animals, a result also observed by Bicalho *et al.* [36] but only during the calving period. Regarding the lactation stage, while Pascottini *et al.* [24] have not highlighted differences in α and β- diversities between the 10^th^ and the 35^th^ DIM, Chen *et al.* [19] obtained significant variations of the Shannon index and Bray-Curtis distances at different AI service times with Holstein cows. Thus, this last study and ours revealed higher Shannon index values as the calving-sampling interval increased. It is logical to think that pregnancy and calving can have a major impact on the vaginal microbiota, which could be caused by the physiological changes or levels of hormones. Indeed, even if Quereda *et al.* [30] only observed differences of beta-diversity between the luteal and follicle phases, they hypothesized that the progesterone level influences the vaginal microbiota composition. This result is in agreement with Laguardia-Nascimiento *et al.* [38], who observed lower α-diversity in pregnant Nellore cows (with high levels of progesterone) compared to non-pregnant animals. Thus, we can hypothesize that the vaginal microbiota could be driven by the host’s levels of hormones, and our new data reinforces this hypothesis.

Overall, the vaginal microbiota appeared to be highly linked with the animal’s physiology and environment but these factors did not explain all the observed variability. The host’s genetics is also known to be a driver of the microbiota composition, for instance in cattle [44] and in the human vagina [45], but this hypothesis has not been investigated in our study.

### Associations between the vaginal microbiota and the productive career length

As previously mentioned, we observed a significant increase of the α-diversity with the number of parities. This result is coherent with an increased diversity according to age, but could also be related with an indirect selection of the animals, with cows showing higher performances and longer lives having a richer microbiota. The productive career is a major issue for breeders, both in terms of cost-efficiency and overall environmental impact [46]. We were not able to highlight any correlation between the α-diversity metrics of the vagina and the career length of the animals. However, we did point out specific associations between the composition of the vaginal microbiota and the longevity of the animals by looking at the global β-diversity and the specific relative abundance of some taxa. Indeed, some genera had been shown to be more abundant in the vagina of long productive career cows, including *Ruminobacter*, *Anaerovibrio*, *Negativibacillus*, *Parasutterella* and two unknown genera from *Paludibacteriaceae* and *Peptococcaceae* families. These genera have already been described in the bovine vagina [30, 47, 48], but little is known about their niches or roles in this reproductive organ. Besides, *Ruminobacter, Anaerovibrio*, *Parasutterella* and *Negativibacillus* have been mostly described in the gut microbiota, where they are generally associated with diverse metabolisms in ruminant species [49–54]. Conversely, *Methylobacterium-Methylorubrum* was generally more abundant in animals with short careers. To our knowledge, *Methylobacterium-Methylorubrum* has not been previously described, neither in the vaginal microbiota of any species, nor in any of the bovine microbiotas. Surprisingly, this genus is a strictly aerobic neutrophil bacterium, while we expect the vagina to be an acidic and closed environment [18]. Maybe its presence was an indicator of the physiological parameters of the vagina. In other studies, this genus was associated with cancer metabolites in humans [55] and appeared to carry antibiotic resistance genes [56]. Interestingly, in addition of being associated with the longevity of the cows, *Negativibacillus* was the only genus observed with a significantly lower abundance in cull animals whereas *Methylobacterium-Methylorubrum* was the second most differentially abundant genus in cull cows out of a total of 49 genera, *Pseudomonas* being the genus with the strongest association with the culling. This genus is known to frequently be resistant to antibiotics [57] and has also been involved in bovine fertility disorders such as endometritis [58] and mastitis [59]. Among the other genera, bacteria considered as pathogenic were observed being more abundant in cull animals, such as *Stenotrophomonas* [60] or *Gallibacterium* [61].

Overall, these results are novel, as our study is the first one to our knowledge, which intended to find associations between the cow longevity and its vaginal microbiota. It will be relevant in the future to evaluate these associations with for instance survival analysis to better appreciate the relationship between the vaginal microbiota and the career length of the dairy cows.

### Associations between vaginal health status and microbiota

As many of the short-career cows carried infection-related bacteria, the associations between the health of the vagina and its microbiota are of major interest in this cohort. However, compared to other studies in cows [24, 25, 62] or sows [15], we did not observe any difference in α or β-diversities between healthy or infected vaginas. A possible explanation could be our loose definition of the infected status, which could have conducted to aggregate multiple infections regardless of the underlying pathogenic agent. Overall, various infectious origins could lead to various kinds of dysbiosis, making them more difficult to group and compare with the healthy vaginas. However, the differential abundance analysis did point out significantly associated genera. Indeed, even if we did not observe some of the typical pathogenic bacteria as *Trueperella* [24], infected vaginas generally had increased abundances of pathogenic genera such as *Fusobacterium*, *Porphyromonas*, and *Peptoniphilus*, which are generally overabundant in the reproductive tracts with metritis [24, 47, 62–64].

Concurrently, other genera were found to be more abundant in healthy animals, *Campylobacter* in particular. Even if adult ruminants exhibit a high amount of *Campylobacter* in their digestive tracts [65, 66], these bacteria are not generally considered as harmless commensal. In a deeper analysis at the ASV level (that is, close to the species taxonomic rank or event strain level), presented the *C. lanienae* as the most significantly overabundant *Campylobacter* species in healthy vaginas. This species is not considered as pathogenic in the literature [67] and its presence could prevent other similar pathogens from being present by establishing on their ecological niche. In this sense, in the digestive tract of beef cattle, the *C. lanienae* has been detected in the large intestine and it has been proposed to prevent the presence of *C. jejuni* [66].

Another intriguing result was the presence of *Streptobacillus*, not well known apart from *S. moniliformis* which was found to be linked to the rat bite fever [68]. Yet, Swartz *et al.* [18] also described it as being an abundant genus of the bovine vagina. Although it remains difficult to conclude on the role of *Streptobacillus* in the vagina because of the lack of knowledge, this genus is part of the Leptotrichiaceae family, as well as *Leptotrichia*, which is known to produce acid, such as lactic acid. Thus, if cows are not sensitive to these two genera, their presence could help to decrease the pH of the vagina and protect it from other pathogens, similarly to *Lactobacillus* in humans [69].

The overabundance of some genera in healthy vaginas was also somehow intriguing as they are generally involved in infections such as *Gallibacterium* in broilers [70]. However, some other findings such as the overabundance of *Lachnospiraceae* UCG-010 was coherent with the previous results of Moreno *et al.* [40] who noticed a similar association between the *Lachnospiraceae* and healthy vaginas. Interestingly, most of the genera significantly overabundant in healthy vaginas could live in anaerobic conditions, meaning that they could better grow in a vagina without oxygen. Thus, maybe these genera did not protect the vagina but were representative of a physiology that prevents contamination by opportunistic aerobic pathogens. This hypothesis would be in agreement with a proposed classification of vaginas depending on the amount of oxygen available for the vaginal microbiota ecosystem [38].

### The vaginal microbiota as a potential indicator of the fertility performances

The fertility of the cows strongly depends on the vaginal health [71]. In contrast to the work of Chen *et al*. [19] which did not find any correlation between the α and β-diversities in the vagina and the pregnancy status of dairy cows, we reported here that both the α and β-diversities were significantly associated with the success of the first insemination (FIS), with reduced diversity being beneficial for conception at first service in Holstein cows. Interestingly, this strong association was reinforced as we observed some genera being overabundant both in healthy vaginas and in animals with enhanced reproductive performances. This is the case of *Leptotrichia* and *Streptobacillus*. *Methanosphaera*, another lactic acid producer, was also associated with the first AI successful outcome and negatively associated with the C-AI_f_ length. Even if *Methanosphaera* is better known for its presence in the digestive tract, this genus has also been found more abundant in pregnant ewes [16]. However, its role and niche in the vagina remains uncertain.

Other genera were also associated with the reproductive performances of the cows. For instance*, Fournierella* was associated with shorter intervals between two successive calvings and between the calving and the following fertilizing AI. *Fournierella,* from the Ruminococcaceae family, is not commonly observed in vagina. Besides, Chen *et al.* [19] who have studied the relationship between the vaginal microbiota and the pregnancy outcome of AI also observed various unknown genera from *Ruminococcaceae* more abundant in bred animals. Thus, deeper analyses need to be performed to better understand the potential roles or niches *Fournierella* has in cows. Besides, apart from *Fournierella*, we also highlighted another genus from the Ruminococacceae family like *Colidextribacter*. Chen *et al.* [19] also introduced unknown genera from the Lachnospiraceae family among the genus that were more abundant in animals that will become pregnant as we did with *Shuttleworthia* from the Lachnospiraceae family. This finding was not surprising as the presence of the Lachnospiraceae family was also considered beneficial by Moreno *et al.* [40].

### The vaginal microbiota is linked with the dairy production performances

Even if the vaginal tract of healthy animals was not expected to directly influence the cows’ production performance, we investigated potential associations between the main dairy traits and the vaginal microbiota. Although we did not evidence correlations between the vaginal α and β-diversities and the dairy performances of the animals, several genera appeared to have a relative abundance that significantly fluctuated with some of the phenotypes of interest: 26, 29 and 26 genera were linked to the milk yield, the fat yield and the protein yield, respectively. In addition, 41 genera had an abundance that was associated with these three phenotypes. Among them, *Lachnospiraceae AC2044 group* was one of the most overabundant in the best performing animals. This genus has not yet been related to the vaginal microbiota, but Liu *et al.* [72] have observed an increased amount of this genus in the rumen microbiota of Holstein cows with higher levels of total milk solid. Conversely, in cows without obvious health problems, we found that animals with lower production performances during the sampled lactation seemed to be have generally a higher relative abundance of pathogenic genera such as *Ureaplasma* [47, 73], *Histophilus* [48] or *Fusobacterium* [24, 47, 48], commonly found in the vagina [30]. Interestingly, as described above we observed that *Fusobacterium was also* more abundant in infected animals, supporting the link between the presence of pathogenic bacteria and reduced milk production. However, as far as we know, no previous studies investigated the association between the vaginal microbiota and milk production traits. Nevertheless, a majority of these differentially abundant genera have been observed in the gut of dairy cows [72], as for the genera which were significantly associated with the productive longevity. As production is often linked to the digestive tract [74, 75], maybe these genera reflected the gut microbiota without being responsible for the differences of performances and were a proxy of the gut microbiota. This last hypothesis has already been brought forward by other studies [37, 38] but calls for further investigations.

### Some vaginal genera were associated with various phenotypes

The present study presented interesting associations between the phenotypes of interest and some bacterial taxa populating the bovine vagina. Some genera seemed to be positively associated with good performances, as for *Negativibacillus* which was more abundant in animals with good production and also increased productive longevity. The presence of this genus in the animals with the best performances was not surprising as it has been involved in digestive mechanisms and good digestive performances [54].

Conversely, other genera had contradictory beneficial effects. These interactions involved traits which are known to be negatively correlated as reproductive and production traits [9]. This was the case for *Leptotrichia* and *Streptobacillus* which were associated with good fertility performances and poorer records in milk, protein and fat yields. Their actions as lactic-acid bacteria could link them with the good fertility records. However, biological associations between their abundance and the production of milk, protein and fat were less evident and have never been reported, even in the digestive tract. Therefore, these bacteria could be biologically associated with only one of the phenotypes and indirectly linked to other traits because of the phenotypic correlations. However, interestingly, some genera which were significantly associated with the fertility traits such as *Fournierella* were not negatively associated with the production traits. In the future, a deeper analysis, as functional analysis with whole metagenome sequencing, which was out of scope for the present study, could help evidence the complex relationships between the vaginal microbiota and the phenotypes.

## Conclusion and perspectives

The microbiota of the dairy cows’ vagina is highly dependent on the cow status and breeding conditions, resulting in a large bacterial α-diversity with a small core microbiota. Associations between this microbiota and several phenotypes are underlined in the present study, considering both the diversity and the composition of the microflora. In addition, we unveiled significant associations between the abundance of some bacterial genera and traits of interest linked to production, reproduction, and health. Overall, our results confirm that the microbiota of this reproductive organ of great interest for the dairy industry should be further studied. Indeed, a better understanding of the close relationship between the vaginal microbiota and its host could offer relevant opportunities to drive this ecosystem and improve the reproductive health status of cows.

## Abbreviations

AI: Artificial insemination
ANCOM-BC: Analysis of compositions of microbiomes with bias correction
ANOVA: Analysis of variance
ASV: Amplicon sequence variant
C-AI_f_: Calving to fertilizing insemination interval
CI: Calving interval
DADA2: Divisive amplicon denoising algorithm 2
FIS: First insemination success
OTU: Operation taxonomic variant
PERMANOVA: Permutational multivariate analysis of variance

## Acknowledgments

The authors acknowledge the breeders and the technicians for the samples and data collection of the study as well as Gérard HEUMEZ (Gènes Diffusion) for its support in the data processing. In addition, the authors thank the INRAE MIGALE bioinformatics facility (MIGALE, INRAE, 2020. Migale bioinformatics Facility, doi: 10.15454/1.5572390655343293E12) for providing the computational resources needed for our analyses.

## Availability of data and materials

The data used in this study are available upon request from the corresponding author.

## Author contributions

CA, SD and GE conceptualized the study and conceived the experimental design. SMa and SMe conducted the whole lab work.

LB performed the bioinformatic data processing and biostatistical analyses under the supervision of JE.

LB, JE, SD, PC, CA, GE, and M-PS all participated in the interpretation of the results.

LB drafted the manuscript.

All authors conceptualized the manuscript, edited later drafts, read, and approved the final manuscript.

## Ethics approval and consent to participate

This work was conducted on farm animals reared for commercial purposes in compliance with the French regulation (*Code Rural et de la Pêche Maritime*) and the European Council Directive 98/58/EC. In accordance with the legislation, no approval from the Institutional Animal Care and Use Committee or ethics committee was necessary as all the performed sampling operations were part of routine animal manipulations by duly authorized technicians. These technicians were employees of the Gènes Diffusion breeding company (Douai, France) who took part in the routine reproduction monitoring. They were holders of the CAFTI diploma (*Certificat d’Aptitude aux Fonctions de Technicien d’Insémination* - Certificate of Fitness for Insemination Technician Functions), approved by the declaration to the EDE (Departmental Establishments of Breeding) that authorizes them for biological sampling in agreement with animal welfare regulation. The farmers involved in this study agreed to the use of their animals’ samples for research purposes.

Our study is reported in full compliance with the ARRIVE guidelines (https://arriveguidelines.org/).

